# Multi-layered Network Analysis of Osteoking in the Treatment of Osteoporosis: Unraveling Mechanisms from Gene Expression to Molecular Docking

**DOI:** 10.1101/2024.03.18.585459

**Authors:** He Chen, Jun Ying, Xianjie Xie, Boyun Huang, Pengcheng Lin

## Abstract

This study aimed to elucidate the therapeutic mechanisms of Osteoking in the treatment of osteoporosis through a comprehensive analysis of potential targets, active ingredients, and associated pathways.

**Method:** The study employed an integrated approach to understand the molecular mechanisms underlying Osteoking’s treatment of osteoporosis. The construction of the protein-protein interaction network involved analyzing data from GENEMANIA and STRING databases. KEGG enrichment analysis was performed to identify enriched pathways, focusing on the cAMP signaling and PI3K-AKT signaling pathways. Active ingredients, disease targets, and osteoporosis-related pathways were integrated into a comprehensive network diagram using Cytoscape.

**Result:** The Gene Expression Omnibus (GEO) database was employed to identify osteoporosis-related gene targets, revealing 3,578 downregulated and 1,204 upregulated genes. Simultaneously, Osteoking’s active ingredients and potential targets were investigated using the Traditional Chinese Medicine Integrated Database (ETCM). A protein-protein interaction network and KEGG pathway enrichment analysis were constructed, highlighting potential targets for Osteoking’s therapeutic effects on osteoporosis. The study also conducted molecular docking analysis, revealing the strong binding capacities of Kaempferol with key disease targets. The results suggest that Osteoking, particularly its active component Kaempferol, holds promise as a potential intervention for osteoporosis, providing insights for further exploration and development of osteoporosis treatments

**Conclusions:** In conclusion, despite some limitations, this study provides valuable information for the treatment of osteoporosis. Future research should make further progress by continually expanding data sources, conducting in-depth experimental validations, and broadening the scope of targets to better understand and address this common skeletal disorder.

**Funding:** This work was supported by the Scientific Foundation of Fuzhou Municipal Health Commission (2021-S-wp3).

## Introduction

Osteoporosis, characterized by diminished bone mass and structural deterioration, stands as a prevailing metabolic skeletal disorder posing a significant global health concern ^[1]^. Its emergence as a major clinical problem is underscored by the escalating aging population, with an anticipated surge in the incidence of osteoporosis-related complications. Despite the prevalent use of conventional pharmaceutical interventions, the associated side effects and suboptimal therapeutic outcomes necessitate exploration of alternative therapeutic approaches ^[2-5]^.

The ancient Chinese medicinal formulation, “Heng Gu Bone Healing Agent” (Osteoking), derived from centuries-old Yunnan Yi ethnic wisdom, has been proposed as a promising intervention for osteoporosis. Documented in the 2020 edition of the “Pharmacopoeia of the People’s Republic of China,” Osteoking amalgamates botanical ingredients such as safflower, notoginseng, ginseng, astragalus, eucommia, honeysuckle, schizophragma, and tangerine peel. Widely employed in traditional Chinese medicine, Osteoking has exhibited efficacy in various bone-related disorders, including fractures, avascular necrosis of the femoral head, lumbar disc herniation, and osteoarthritis^[6-8]^.

While modern clinical studies have hinted at Osteoking’s potential to promote calcium deposition, enhance bone mass, and restore trabecular architecture, its complex composition and multifaceted mechanisms have impeded widespread clinical adoption. Hence, unraveling the intricate network relationships among drugs, targets, genes, and diseases becomes imperative to elucidate Osteoking’s pharmacological basis for mitigating osteoporosis.

This study employs an integrative approach, incorporating gene expression analysis, network pharmacology, and molecular docking, to comprehensively investigate the potential mechanisms underpinning Osteoking’s efficacy in osteoporosis treatment. Through the exploration of drug-target interactions, gene-pathway associations, and molecular docking validations, we aim to shed light on the nuanced pharmacological actions of Osteoking, providing valuable insights for its clinical application in the context of osteoporosis management.

## MATERIALS AND METHODS

### Osteoporosis Target Collection

The Gene Expression Omnibus (GEO) database (www.ncbi.nlm.nih.gov/geo/) was utilized to retrieve disease gene targets associated with osteoporosis. A total of 4 samples from normal individuals and 4 samples from osteoporosis patients were extracted from the GSE35959 dataset. Differential gene analysis was performed using GEO analysis tools, and volcano plots and differential heatmaps were generated using ggplot2 and pheatmap packages in R software to identify osteoporosis-related targets.

### Osteoking Active Ingredient and Target Collection

The Traditional Chinese Medicine Integrated Database (ETCM, http://www.tcmip.cn/ETCM/index.php/Home/Index/) was employed to search for active ingredients and potential targets of Osteoking ^[9]^. Using keywords such as Chenpi (CITRI RETICULATAE PERICARPIUM), Duzhong (EUCOMMIAE CORTEX), Honghua (CARTHAMI FLOS), Huangqi (ASTRAGALI RADIX), Renshen (GINSENG RADIX ET RHIZOMA), Sanqi (NOTOGINSENG RADIX ET RHIZOMA), Yangjinhua (DATURAEFLOS), and Zuandifeng (SCHIZOPHRAGMA INTEGRIFOLIUM) in Chinese Pinyin for retrieval purposes, the corresponding academic terminologies were saved to document information on the active ingredients and targets for each drug.

### Protein-Protein Interaction Network Construction

The Venny website (https://bioinfogp.cnb.csic.es/tools/venny/index.html) was utilized to identify the intersection between osteoporosis-related targets and Osteoking drug targets. The overlapping targets were considered potential therapeutic targets of Osteoking for osteoporosis. The targets were then imported into the GENEMANIA website (http://genemania.org) to establish their associations ^[10]^. Additionally, the targets were input into the STRING database, and the resulting interaction data were exported in CSV format for constructing a protein-protein interaction network using Cytoscape software ^[11]^. Topological analysis was conducted to assess the degree centrality of each node in the network.

### KEGG Pathway Analysis

The intersection of targets was subjected to KEGG pathway enrichment analysis (http://bioinfo.org/kobas/) ^[12]^. The results were visualized using R software to generate bubble plots. Relevant pathways associated with osteoporosis were selected from the enrichment results and visualized in a bubble plot. The enriched pathway-related targets were used to construct a network diagram in Cytoscape, representing the relationships among drugs, active ingredients, targets, and pathways.

### Molecular Docking

Based on topological analysis results, the 3D structures of active ingredients and targets were obtained from the PubChem and PDB databases. Using CB-DOCK with the Number of cavities for docking set to 5, docking simulations were performed, and the results were visualized using PyMOL (The PyMOL Molecular Graphics System, Version 2.0 Schrödinger, LLC.) for further analysis of the docking interactions ^[13]^.

## RESULT

### Acquisition of Osteoporosis Disease Targets

Data analysis of GSE35959 from the Gene Expression Omnibus (GEO, www.ncbi.nlm.nih.gov/geo/) was performed to identify disease-related gene targets in osteoporosis. Using GEO2R with predefined parameters for P-value and LogFC, the gene expression differences in bone marrow mesenchymal stem cells between osteoporosis patients and healthy individuals were analyzed. The differential gene analysis revealed a total of 3,578 downregulated genes and 1,204 upregulated genes (Figure 1A). Subsequently, these differentially expressed genes were visualized through hierarchical clustering, illustrating a distinct separation between osteoporosis patients and normal individuals (Figure 1B).

**Figure 1:**
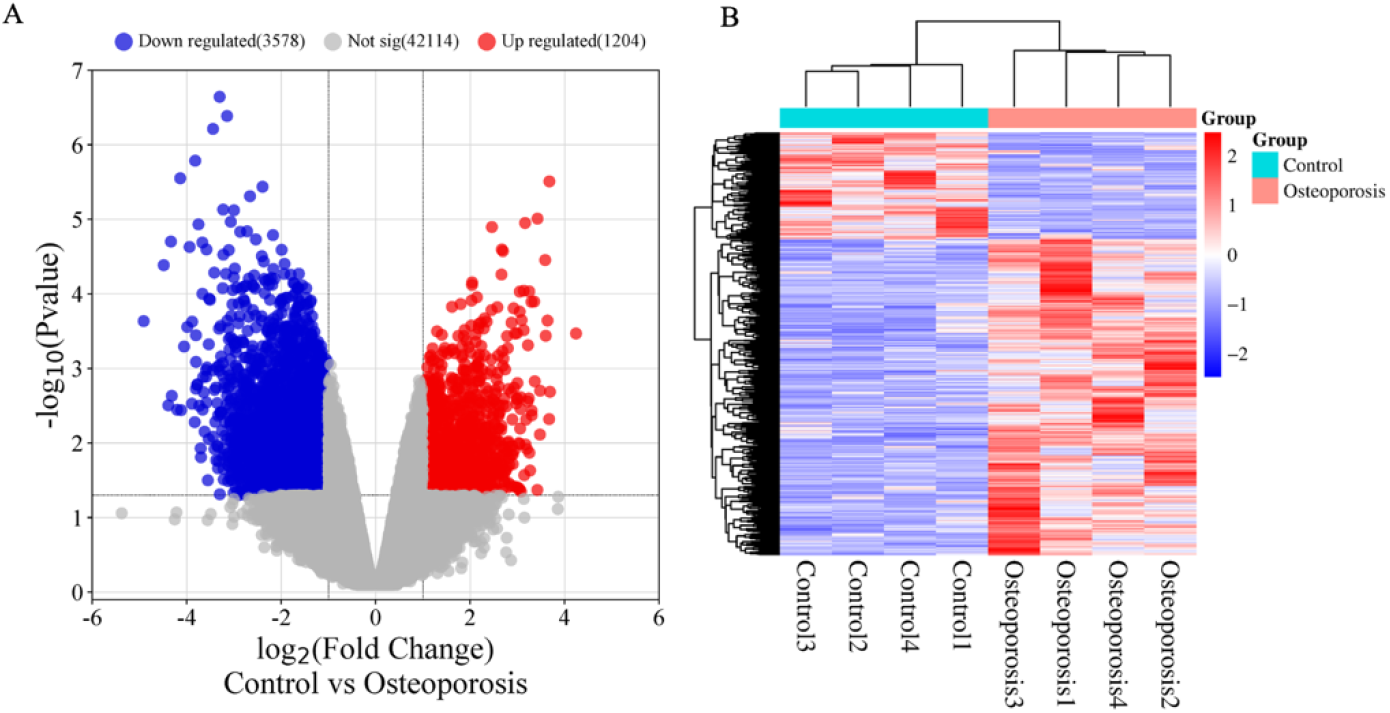
(A) Display of volcano maps filtered using P and FC values. (B) Cluster heatmap of differentially expressed genes.

### Identification of Osteoking Drug Targets

In the ETCM database (http://www.tcmip.cn/ETCM/index.php/Home/Index/), the names of each drug were entered to identify their potential active components and associated targets. A correlation coefficient greater than 0.8 was considered an acceptable threshold for potential therapeutic targets. The target information was exported as a CSV file for each individual drug, and after merging and removing duplicates, a total of 433 targets were recognized as potential targets for Osteoking.

### Construction of Drug-Disease Target Network

Utilizing the Venny online tool (https://bioinfogp.cnb.csic.es/tools/venny/index.html), an analysis was conducted to determine the intersection of osteoporosis disease targets and Osteoking drug targets, resulting in 57 common targets (Figure 2A). These targets were regarded as potential active targets for Osteoking in treating osteoporosis. The drug ownership of these targets and the intersection with disease targets were imported into Cytoscape for topological analysis (Figure 2B). GINSENG RADIX ET RHIZOMA had the highest degree (44), followed by ASTRAGALI RADIX with a degree of 41, and then CARTHAMI FLOS with a degree of 33, suggesting varying degrees of importance in the Osteoking treatment network (Figure 2C).

**Figure 2:**
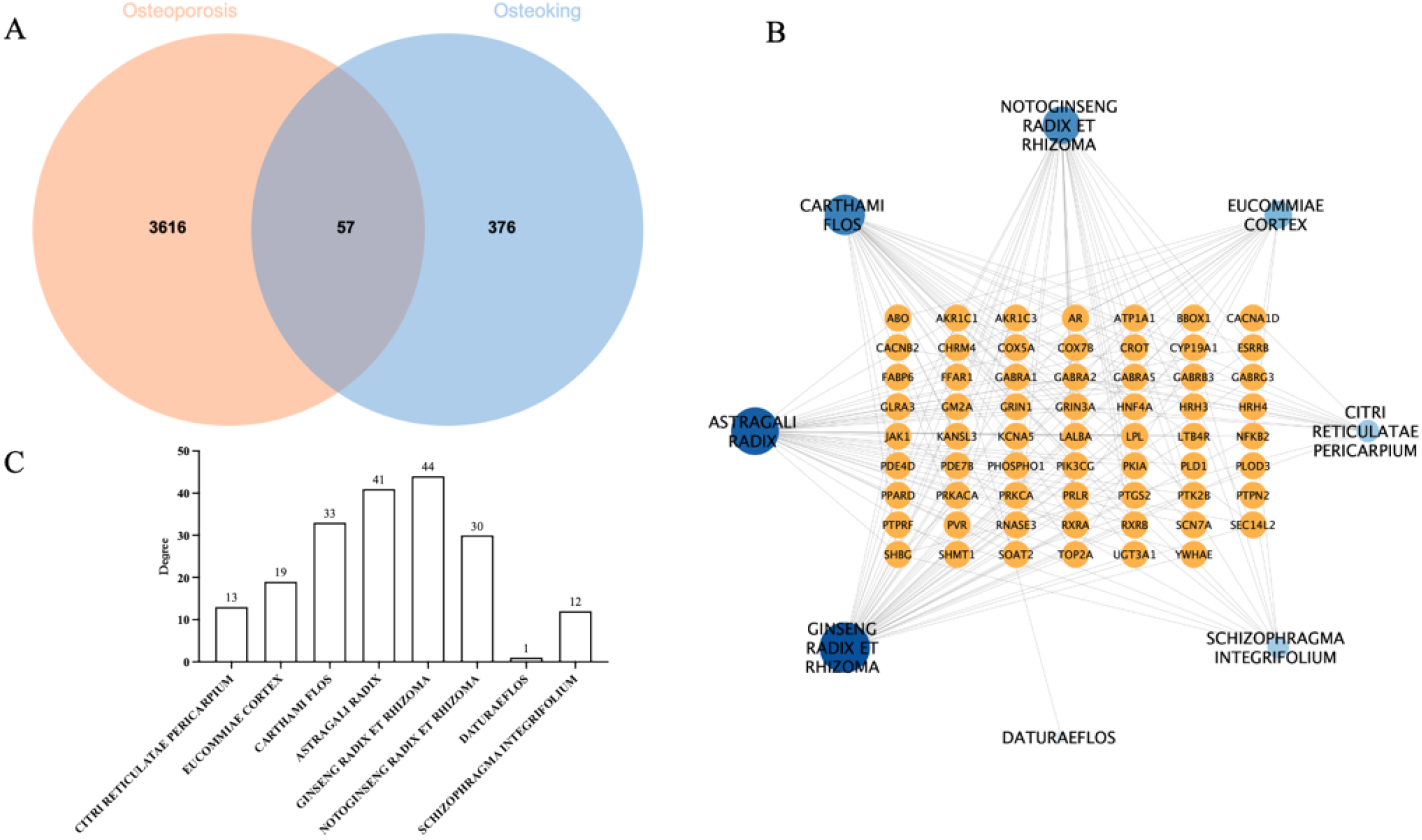
(A) Intersection of Veeny plots of Osteoporosis disease targets and Osteoking drug targets. (B) Osteoking drug - Osteoporosis target network diagram. (C) Network topology analysis results.

### PPI Network Construction

The potential active targets of Osteoking for osteoporosis treatment were imported into the GENEMANIA website to explore their interactions. The analysis revealed that within the 57 common targets, co-expression constituted 35.96%, physical interactions 22.04%, predicted 13.38%, pathway 9.45%, genetic interactions 7.26%, shared protein domains 6.85%, and co-localization 5.05% (Figure 3A). Subsequently, STRING database analysis was performed to further assess the interaction relationships among targets. Topological analysis using Cytoscape, which included Degree, Betweenness Centrality, and Closeness Centrality data (Figure 3B-C), along with network analysis using CytoHubba (Figure 3D), identified core targets such as PRKACA, PTGS2, and PRKCA, suggesting their crucial role in Osteoking’s effectiveness in treating osteoporosis.

**Figure 3:**
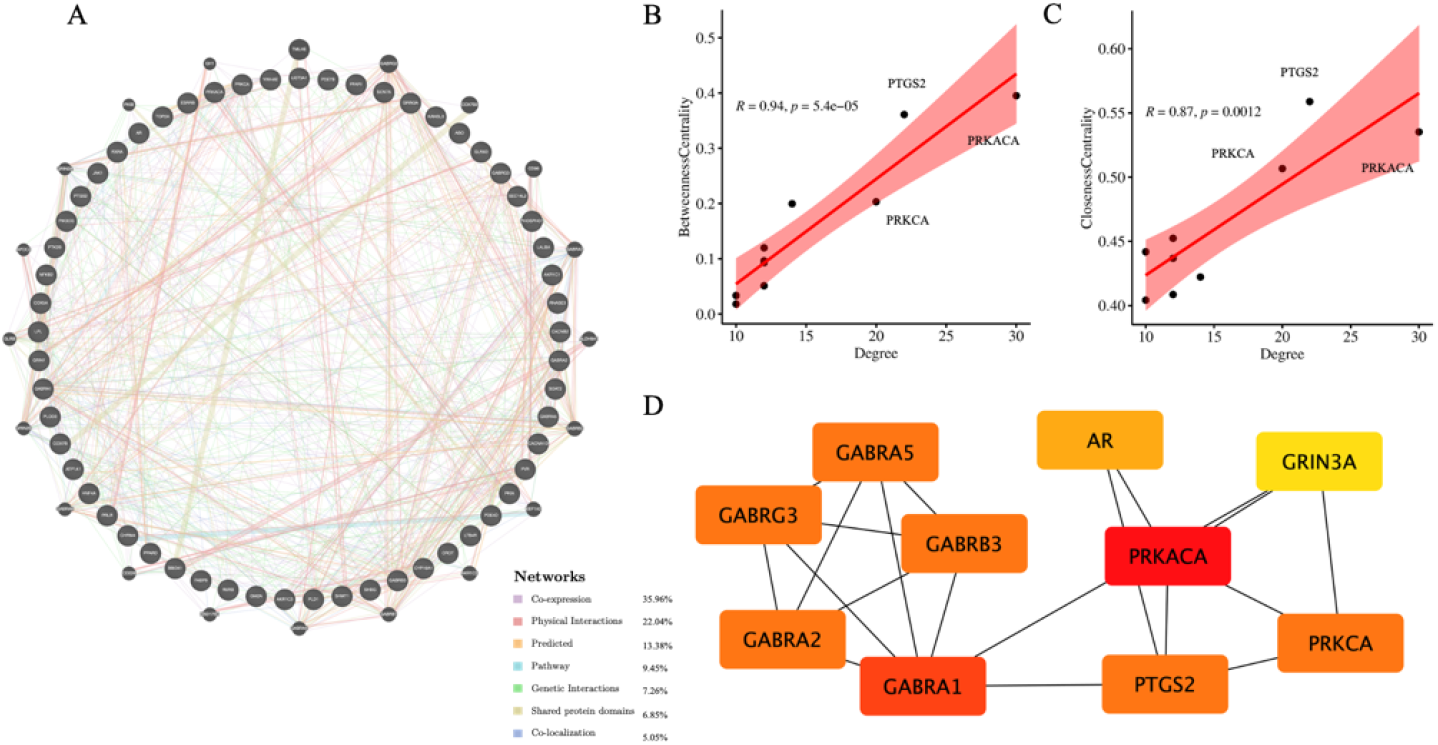
(A) Network diagram of interactions between targets. (B) Correlation analysis of the degree, betweenness centrality, and closeness centrality of the interaction network between Osteoporosis and Osteoking targets in topological analysis. (C) Visualization processing of core target results obtained by CytoHubba.

### KEGG Enrichment Analysis

To explore the potential mechanisms and actions of Osteoking in osteoporosis, the 57 targets were subjected to KEGG pathway enrichment analysis. The enrichment results revealed significant pathways, including the cAMP signaling pathway and PI3K-AKT signaling pathway (Figure 4A). Further exploration of osteoporosis-related pathways identified by P<0.05 included Parathyroid hormone synthesis, secretion and action, Insulin secretion, PI3K-Akt signaling pathway, Calcium signaling pathway, MAPK signaling pathway, Wnt signaling pathway, Osteoclast differentiation (Figure 4B).

**Figure 4:**
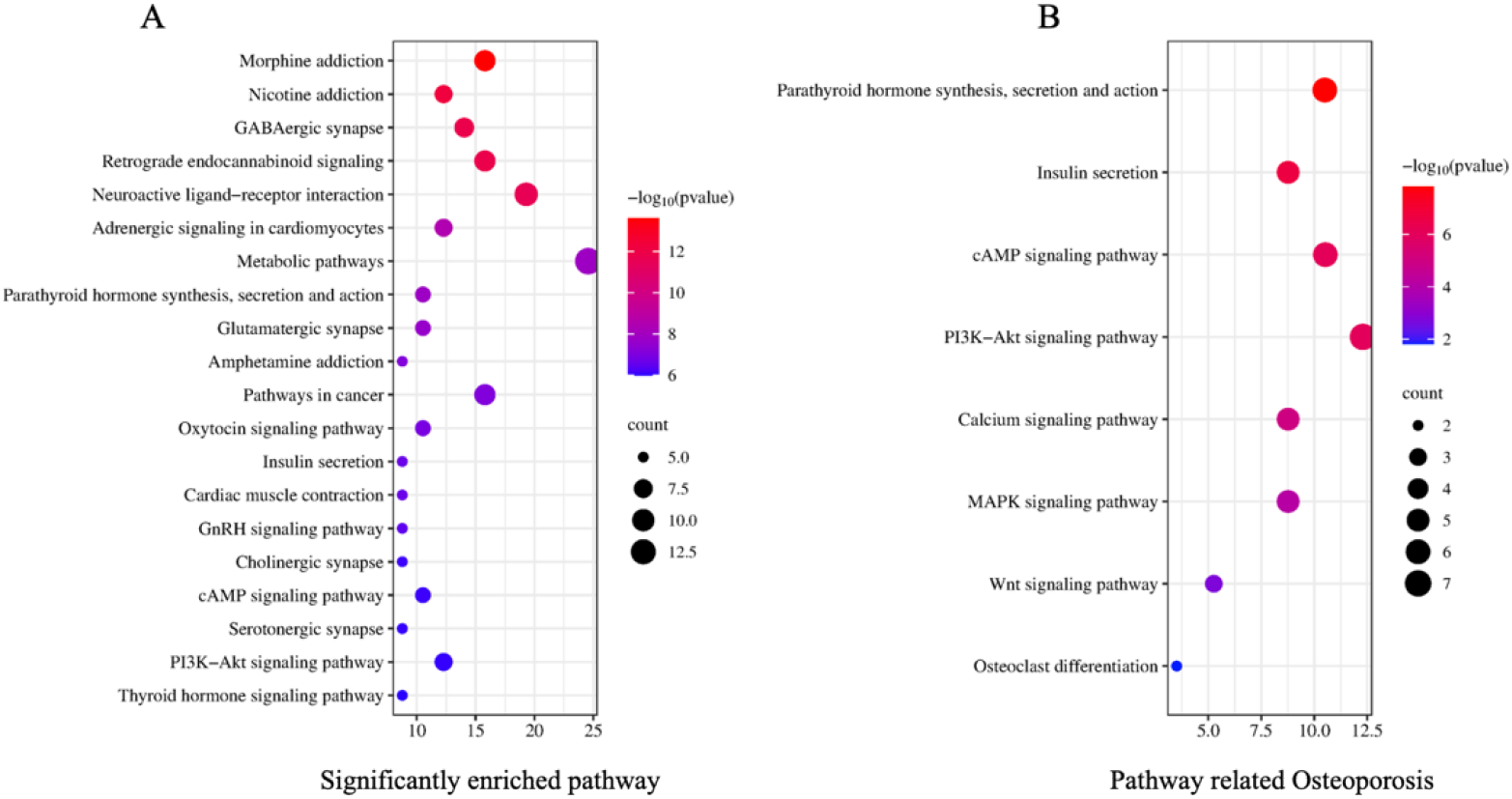
(A) KEGG enrichment analysis of drug disease target intersection. (B) Osteoporosis related KEGG analysis.

### Active Ingredient-Disease Target Network

Utilizing the potential active ingredients obtained from the ETCM database, we collected the targets associated with these active ingredients. The visualization of the active ingredient-disease target network was performed in Cytoscape (Figure 5A). Simultaneously, a topological analysis of the potential active substances in Osteoking’s action against osteoporosis was conducted. In the active ingredient network, Cetylic Acid, Sitosterol, and Kaempferol were identified as the top three nodes based on degree centrality in the network topology analysis. These components may represent crucial substance foundations for Osteoking’s efficacy in treating osteoporosis, with the potential targets being PRKACA, AR, and GABRA1, indicating their close involvement in the pharmacological relationship.

**Figure 5:**
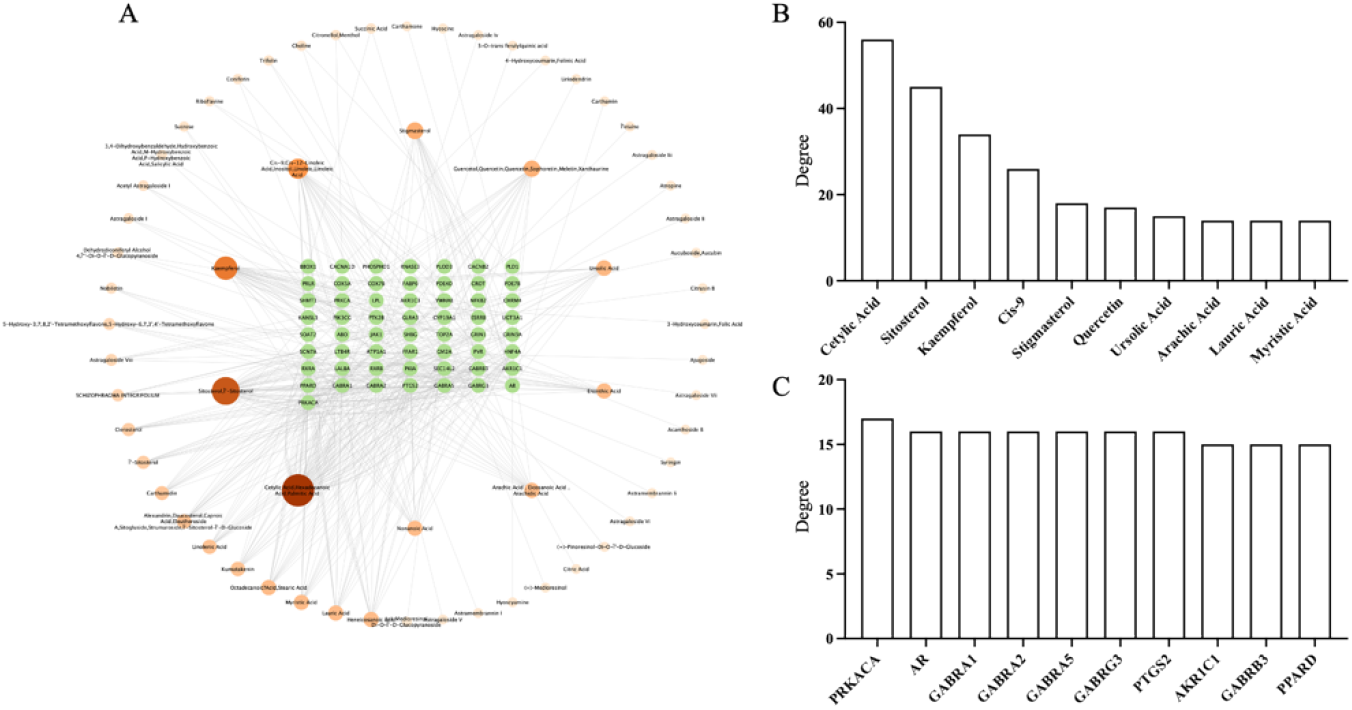
(A) Osteoking drugs-potential active ingredients-disease target network. (B-C) Network topology analysis results.

### Drug-Active Ingredient-Disease Target-Disease-Associated Pathway Network

By integrating the previously enriched targets from osteoporosis-related pathways, along with drugs, active ingredients, and targets, a multi-layered network diagram was constructed using specialized software (Figure 6A). This comprehensive analysis aimed to elucidate the potential mechanisms through which the drug may exert its therapeutic effects on the disease. The analysis revealed that the drugs Cetrylic Acid, Cis-9, and Kaempferol played crucial roles in Osteoking’s treatment of osteoporosis. Additionally, disease targets PRKACA, RXRA, and PPARD were identified as pivotal nodes in this network, underscoring their significant contributions to the therapeutic process (Figure 6B-C).

**Figure 6:**
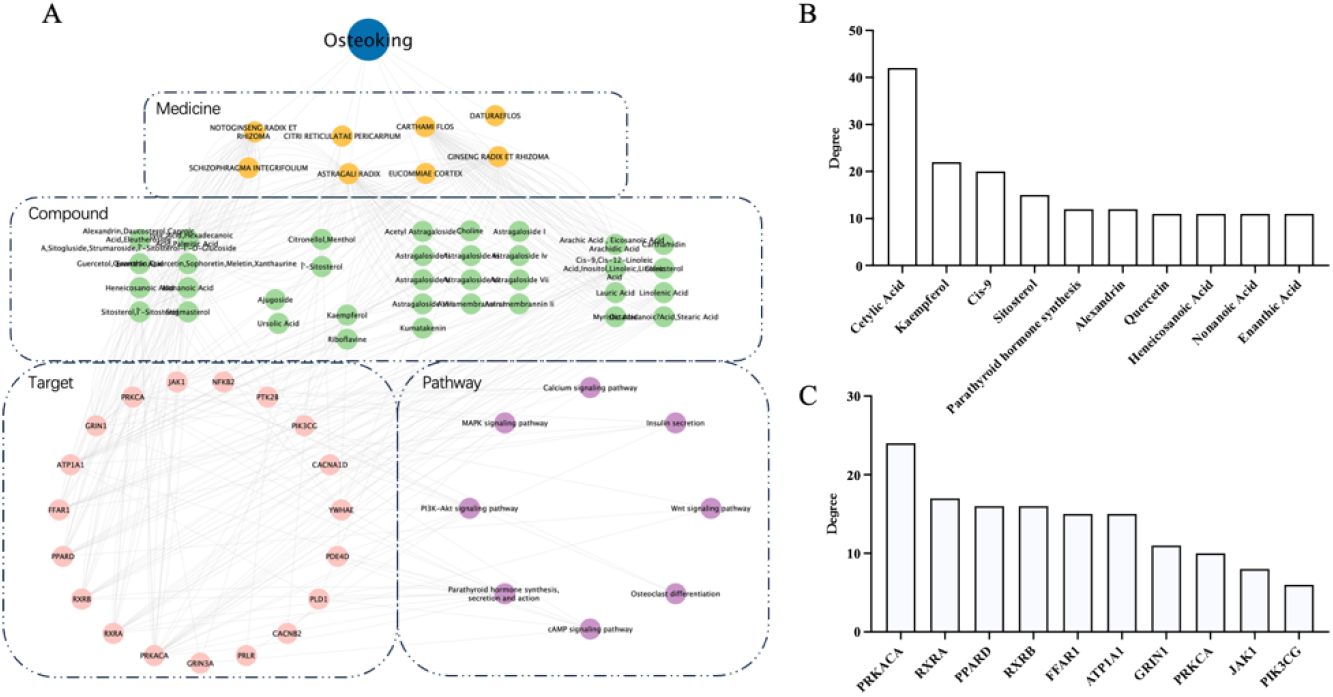
(A) Osteoking drugs-potential active ingredients-disease targets-pathway network diagram. (B-C) Network topology analysis results.

### Molecular Docking Results

The 3D structures of Cetrylic Acid, Cis-9, and Kaempferol were obtained from the PubChem database, while the structures of disease targets PRKACA (PDB ID:2AWH), RXRA (PDB ID:1BY4), and PPARD (PDB ID:2GU8) were acquired from the PDB website. Virtual docking of small molecule drugs and large molecule proteins was conducted using CB-DOCK2, and the results were visualized using PyMOL software. Molecular docking analysis revealed favorable binding affinities between small molecule drugs and disease targets, the binding energy is all less than 5kcal/mol. Notably, Kaempferol exhibited the strongest binding capacities with PRKACA, RXRA, and PPARD, with binding energies of −8.7 kcal/mol, −9.0 kcal/mol, and −8.4 kcal/mol, respectively. These findings suggest that Kaempferol may be a crucial component of Osteoking in the treatment of osteoporosis (Figure 7).

**Figure 7:**
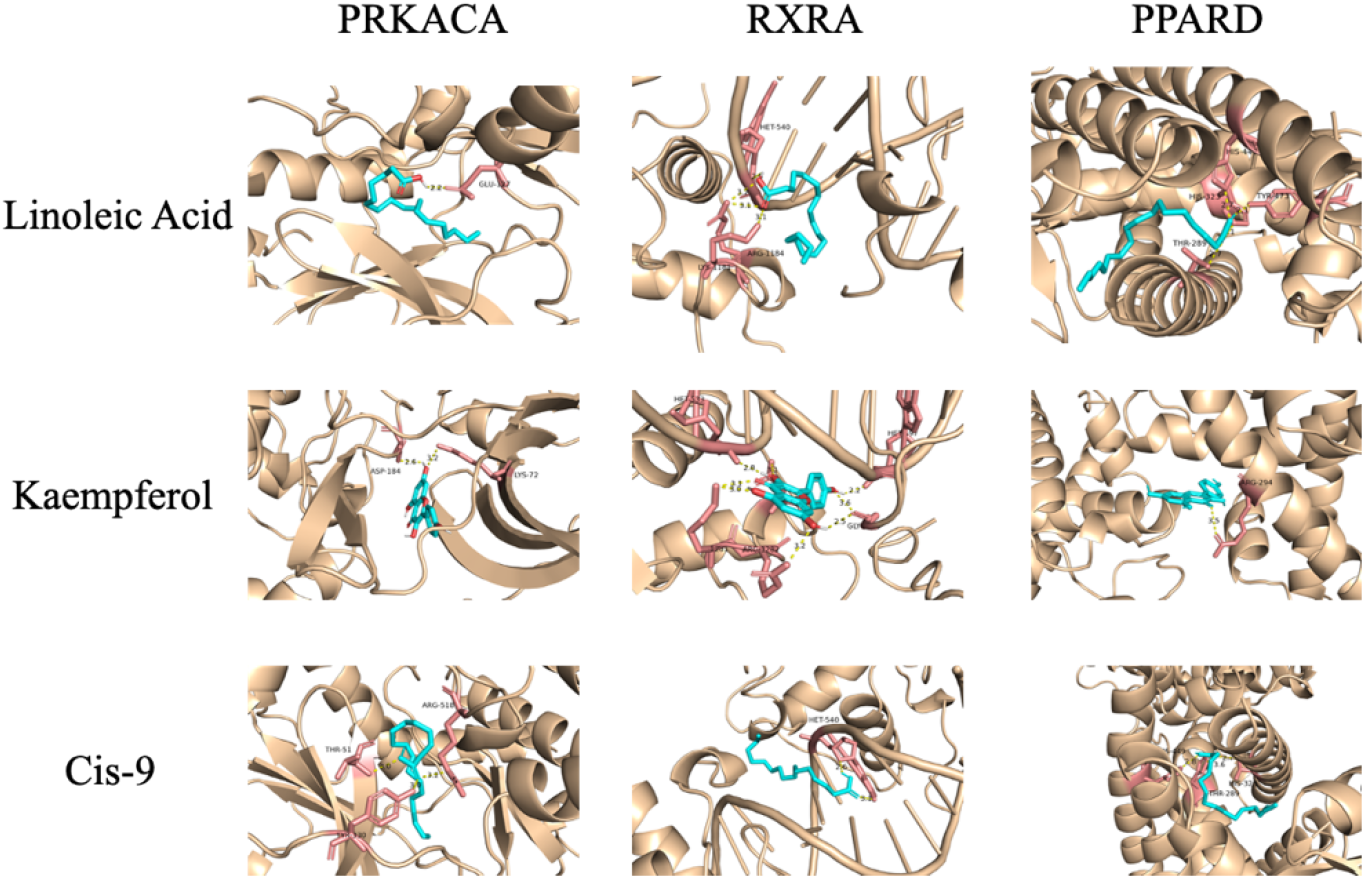
Results of the key chemical components of Osteoking molecular docking.

## DISCUSSION

Osteoporosis, as a globally prevalent chronic malady, exerts a profound impact on patients’ quality of life and imposes substantial burdens upon both healthcare systems and socioeconomic structures^[14]^. The primary objective of this investigation is to gain a comprehensive understanding of the molecular mechanisms underpinning osteoporosis. Through a holistic analysis encompassing the Gene Expression Omnibus (GEO) database and the pharmaceutical agent Osteoking, this study has yielded novel insights into various facets of this pathological condition.

Primarily, employing data mining techniques on the GEO database facilitated the identification of a plethora of differentially expressed genes within the GSE35959 dataset, including both upregulated and downregulated genes. This outcome not only contributes novel insights into the genetic underpinnings of osteoporosis but also establishes a foundational basis for future investigations seeking innovative therapeutic targets. The discerned alterations in the gene expression profile of bone marrow mesenchymal stem cells under pathological conditions emphasize pivotal clues for comprehending the molecular regulatory mechanisms governing osteoporosis.

Subsequently, through an in-depth scrutiny of the Osteoking pharmaceutical, we have unveiled its potential therapeutic mechanisms against osteoporosis. The pharmaceutical composition of Osteoking encompasses diverse active constituents intricately linked to pertinent pathways and biological processes associated with osteoporosis. The construction of an active ingredient-disease target network and a protein-protein interaction network has identified specific components, such as GINSENG RADIX ET RHIZOMA and ASTRAGALI RADIX, playing crucial roles within the overall treatment network. This not only furnishes the molecular foundation for Osteoking’s efficacy in osteoporosis treatment but also guides future endeavors towards the development of more precision-oriented therapeutic strategies. These findings carry profound clinical significance. Primarily, a nuanced understanding of the molecular mechanisms underlying osteoporosis facilitates enhanced prediction of disease risk among patients, enabling timely intervention and personalized treatment modalities. Secondly, an in-depth comprehension of Osteoking contributes to the optimization of its treatment protocols, thereby augmenting its efficacy among osteoporotic patients. This holds paramount value in enhancing patient quality of life and alleviating the burdens on healthcare resources.

Our identification of 3578 downregulated genes and 1204 upregulated genes within the GSE35959 dataset reflects crucial involvement of bone marrow mesenchymal stem cells in the onset of osteoporotic symptoms. The increased abundance of downregulated genes may be associated with diminished functionality of patients’ bone marrow mesenchymal stem cells and a decline in the capacity for bone formation and repair. The emergence of upregulated genes suggests the activation of biological processes such as inflammation and cellular metabolism abnormalities, playing pivotal roles in the development of osteoporosis. The discovery of differentially expressed genes profoundly reveals the biological distinctions between osteoporotic patients and normal individuals. Under normal circumstances, the gene expression pattern of bone marrow mesenchymal stem cells typically maintains the structure and functionality of the skeleton. Among the upregulated genes, pathways involving inflammation, bone resorption, and lipid metabolism were implicated, closely aligning with known pathogenic mechanisms of osteoporosis. Simultaneously, the enrichment of downregulated genes pertained to bone marrow mesenchymal stem cell differentiation, bone cell formation, and bone matrix synthesis, suggesting that the abnormal functionality of bone marrow stem cells in osteoporotic patients may be a crucial link in the disease’s progression. These findings bear significant clinical implications.

As a representative of traditional Chinese medicine formulations, the selection of Osteoking’s pharmaceutical and active ingredients is well-considered, based on a profound understanding of the extensive history and widespread use of Chinese herbs in the treatment of osteoporosis. This rational selection not only reflects the accumulated knowledge of traditional medicine in the field of osteoporosis research but also provides robust support for exploring novel treatment strategies. We delved into the historical and current status of key components in Osteoking. Ingredients such as GINSENG RADIX ET RHIZOMA, ASTRAGALI RADIX, and CARTHAMI FLOS have long been considered in traditional Chinese medicine to have the efficacy of nourishing qi and blood, strengthening tendons and bones ^[15-20]^. Through an analysis of the active ingredient-disease target network and topological analysis, we emphasize the crucial roles of components such as GINSENG RADIX ET RHIZOMA, ASTRAGALI RADIX, and CARTHAMI FLOS in the Osteoking network for treating osteoporosis. These components were found to have high degree values in the network, suggesting their potential significance throughout the treatment process. GINSENG RADIX ET RHIZOMA, as the component with the highest degree value in the network, has been widely used in traditional Chinese medicine for invigorating the body and nourishing qi and blood. ASTRAGALI RADIX and CARTHAMI FLOS are well-known in clinical practice for their properties of replenishing qi and promoting blood circulation ^[21, 22]^. These components may impact the pathogenic mechanisms of osteoporosis through multiple pathways, including promoting bone cell proliferation, inhibiting bone resorption, and improving the functionality of bone marrow mesenchymal stem cells ^[23, 24]^. Further comparison of our findings with existing literature reveals consistency in the selection of Osteoking’s components with other research results ^[25, 26]^. By corroborating our experimental results, we further consolidate the feasibility and potential efficacy of Osteoking in the treatment of osteoporosis.

The construction of protein-protein interaction networks is a crucial aspect in the investigation of the therapeutic mechanisms of osteoporosis. In our study, we successfully delineated the protein-protein interaction network of Osteoking’s therapeutic effects on osteoporosis by integrating the analysis results from the GENEMANIA and STRING databases. This network not only provides a comprehensive perspective on understanding the mechanism of action of Osteoking but also underscores the significance of its core targets, such as PRKACA, PTGS2, and PRKCA. Firstly, it is essential to emphasize that the construction of protein-protein interaction networks is not merely a presentation of molecular-level interactions but also a grasp of potential key nodes throughout the entire therapeutic process. The high connectivity of core targets like PRKACA, PTGS2, and PRKCA in the network implies their potential critical roles in Osteoking’s treatment of osteoporosis. PRKACA has been identified as significantly associated with skeletal disorders, with mutations in PRKACA being notably prevalent in individuals with skeletal defects. Furthermore, its correlation with the proliferation of osteoblasts underscores its pivotal role in bone health ^[27, 28]^. PTGS2 is a key protein regulating inflammatory responses and cellular apoptosis, and its modulation in bone metabolism may be related to Osteoking’s anti-inflammatory and pro-survival effects on bone cells ^[29-31]^. PRKCA, a protein kinase C isoform, is intricately linked to crucial biological processes such as cell proliferation, differentiation, and apoptosis ^[32, 33]^. The identification of these core targets holds promise for providing clues towards unraveling the molecular mechanisms underlying Osteoking’s treatment of osteoporosis.

KEGG enrichment analysis represents a critical step in our study, as its results not only offer a series of enriched pathway mechanisms but also deepen our understanding of the potential effects of Osteoking on treating osteoporosis. In our investigation, pathways such as the cAMP signaling pathway and PI3K-AKT signaling pathway were found to occupy significant positions in the enrichment results. These pathways have been extensively studied in the pathophysiological processes of osteoporosis and are closely associated with the regulation of bone cell proliferation, differentiation, and metabolism. The cAMP signaling pathway plays a crucial role in bone formation by activating key molecules like cAMP response element-binding protein (CREB), regulating the expression of various genes, thus influencing the functionality of bone cells ^[34, 35]^. On the other hand, the PI3K-AKT signaling pathway plays a pivotal role in regulating crucial biological processes such as cell survival, proliferation, and differentiation, exerting a significant impact on the activity of bone cells and bone formation ^[36, 37]^. The discovery of these enriched pathway mechanisms provides crucial clues for explaining the effects of Osteoking in treating osteoporosis. Firstly, the enrichment of the cAMP signaling pathway may suggest that Osteoking modulates intracellular cAMP levels, affecting the expression of relevant genes, thereby mediating the regulation of bone cell proliferation, differentiation, and bone matrix generation. Secondly, the enrichment of the PI3K-AKT signaling pathway indicates that Osteoking may promote the survival and proliferation of bone cells by activating the PI3K-AKT signaling pathway, thereby aiding in the repair and regeneration of bone tissue.

Integrating active ingredients, disease targets, and osteoporosis-related pathways into a comprehensive network diagram is a crucial step in our study. This not only provides a deeper insight into the understanding of the treatment mechanism but also highlights the key roles of active ingredients such as Cetylic Acid, Sitosterol, and Kaempferol in the entire treatment network. Firstly, the integration of the active ingredient-disease target network emphasizes the potential mechanisms of action of these active ingredients in treating osteoporosis. Through Cytoscape visualization, we clearly observe the associations between Cetylic Acid, Sitosterol, and Kaempferol with disease targets like PRKACA, RXRA, PPARD. This underscores that these active ingredients may influence biological processes related to osteoporosis by regulating these key targets. Secondly, the integrated network diagram highlights the comprehensive roles of these active ingredients throughout the entire treatment network. Taking Cetylic Acid as an example, its close connection with PRKACA in the active ingredient-disease target network aligns with its crucial role in the cAMP signaling pathway implicated in KEGG enrichment analysis. At the same time, we tested the good binding ability of these targets and active ingredients through molecular docking. This multi-level association suggests that these active ingredients may exert comprehensive regulation on osteoporosis by modulating multiple pathways and targets.

Despite achieving encouraging results in this study, it is important to acknowledge some limitations. Firstly, the data primarily relies on public databases such as GEO, ETCM, GENEMANIA, and STRING. While these databases provide rich information, they are subject to inherent limitations, such as sample size and heterogeneity in experimental conditions. Future research can enhance the reliability of study results by incorporating more diverse samples and finer experimental designs. Secondly, although molecular docking results suggest that Kaempferol may play a crucial role in Osteoking’s treatment, laboratory results are preliminary and limited to in vitro data. To comprehensively understand the clinical application potential of Kaempferol, more experiments, even validation in animal models, are needed. This will help determine the pharmacokinetics and toxicity characteristics of Kaempferol, providing support for its further development as a potential therapeutic drug. Future research directions should include in-depth investigations into other potential targets. While this study focused on key targets such as PRKACA, RXRA, and PPARD, osteoporosis is a complex disease involving multiple signaling pathways and biological processes. Exploring more targets, especially those closely related to the pathogenic mechanisms of osteoporosis, will help enhance the comprehensiveness and effectiveness of treatment.

## CONCLUSION

In conclusion, despite some limitations, this study provides valuable information for the treatment of osteoporosis. Future research should make further progress by continually expanding data sources, conducting in-depth experimental validations, and broadening the scope of targets to better understand and address this common skeletal disorder.

